# Design and synthesis of DNA origami nanostructures to control TNF receptor activation

**DOI:** 10.1101/2024.04.27.591448

**Authors:** Göktuğ Aba, Ferenc A. Scheeren, Thomas H. Sharp

**Affiliations:** Department of Cell and Chemical Biology, Leiden University Medical Center, 2300 RC Leiden, The Netherlands

**Keywords:** DNA nanotechnology, Tumour necrosis factor receptors, TNF-related apoptosis-inducing ligand (TRAIL), Death receptor 5 (DR5), Receptor clustering

## Abstract

Clustering of type II tumour necrosis factor (TNF) receptors (TNFRs) is essential for their activation, yet currently available drugs fail to activate signalling. Some strategies aim to cluster TNFR by using multivalent streptavidin or scaffolds based on dextran or graphene. However, these strategies do not allow for control of the valency or spatial organisation of the ligands, and consequently control of the TNFR activation is not optimal. DNA origami nanostructures allow nanometre-precise control of the spatial organization of molecules and complexes, with defined spacing, number and valency. Here, we demonstrate the design and characterisation of a DNA origami nanostructure that can be decorated with an engineered single-chain TNF-related apoptosis-inducing ligand (SC-TRAIL) complexes, which show increased cell killing compared to SC-TRAIL alone on Jurkat cells. The information in this chapter can be used as a basis to decorate DNA origami nanostructures with various proteins, complexes or other biomolecules.

## 1 Introduction

Cells sense extracellular signals and change their cellular functions in response.[1,2] Oftentimes, selective receptor-ligand interactions are the initial step for cellular activation.[1,3] In particular, clustering of cell surface receptors is essential for activation of Tumor necrosis factor (TNF) receptor superfamily (TNFRSF), which has important roles in mammalian cell proliferation, cell death, immune regulation and morphogenesis.[4-6] The spatial organization of the cell surface receptors can be crucial for the regulation of cellular signaling cascades. This has been described for a number of receptor-ligand interactions, for example TNF-related apoptosis-inducing ligand (TRAIL) and death receptor 5 (DR5).[1,7-9]

Despite the current knowledge about the structure of the TNFRSF receptors (TNFRs), currently available drugs fail to comprehensively activate signaling in type II TNFRs, such as 4-1BB, CD40 and death receptor 5 (DR5).[10,11] Interestingly, efficient receptor activation was shown when their ligands were presented in membrane-associated form (**Figure 1A**).[11,12] Many agonistic antibodies require crosslinking for effective induction of the receptors.[4,13] Furthermore, the interligand distance affects signaling activation in immunological synapses.[14,15] Some strategies multimerize ligands using multivalent streptavidin, or scaffolds based on peptides, dextran, or graphene, to successfully initiate DR5-mediated cell signaling (**Figure 1B**).[16-21] However, these methods do not allow precise control over the interligand distance, and achieving a ligand arrangement with sub-10 nm distancing is challenging.[15,22] In comparison, DNA origami nanostructures[15,23,24] are an ideal alternative; DNA origami nanostructures allow nanometer-precise control over the spatial organization of ligands, with defined composition and number (**Figure 2**).[25] Moreover, DNA nanostructures are stable in the presence of human sera.[26] DNA nanostructures have been previously used to activate Toll-like receptors using CpG-motifs[25], DR5 using TRAIL-like peptides[15] and T cell antigen receptors (TCRs) by peptide/major histocompatibility complexes (pMHCs).[27] We will not cover all terminology of DNA nanotechnology, but several excellent reviews are available as resources (see e.g.,[28]).

**Figure 1.**
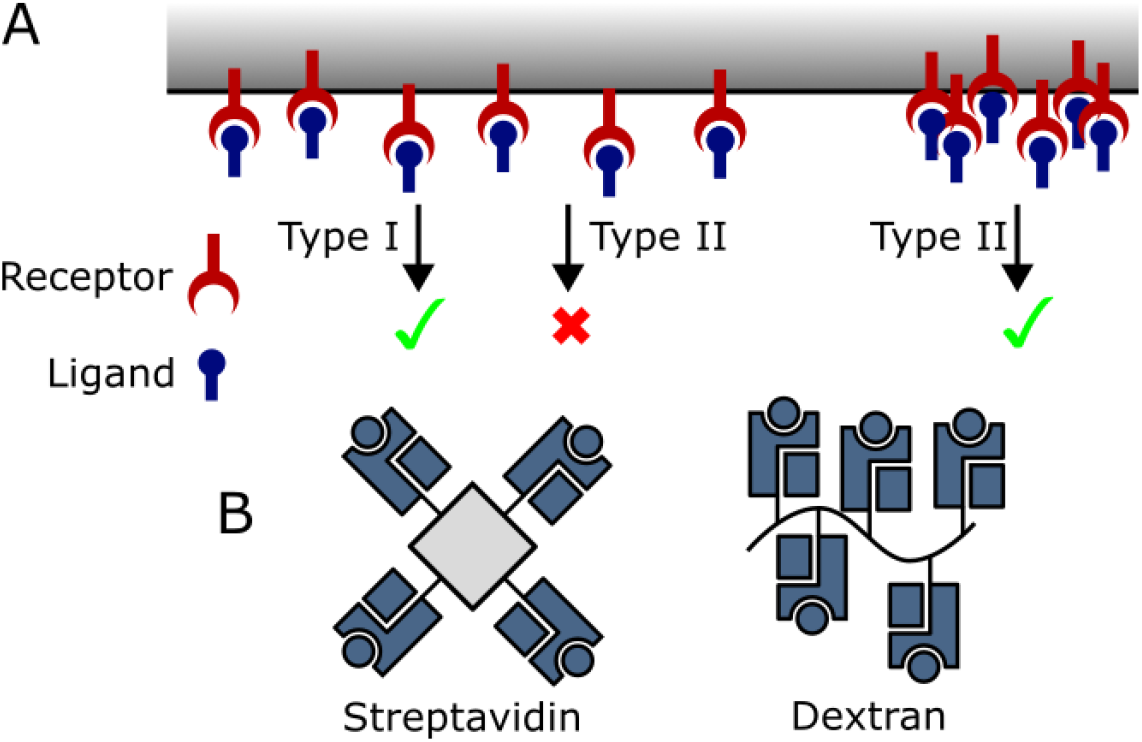
Receptor clustering leads to activation of TNFR. **A)** Schematic overview of type I and type II TNFRs. Type II TNFRs require receptor clustering for activation of signalling, unlike type I TNFRs, which can be activated by the interaction between the ligand and the receptor.[11,12] **B)** Schematic representation of two different multimerization techniques of major histocompatibility complexes (MHCs). MHCs are tetramerized using streptavidin on the left and multimerized on a dextran backbone on the right.[20,21]

**Figure 2.**
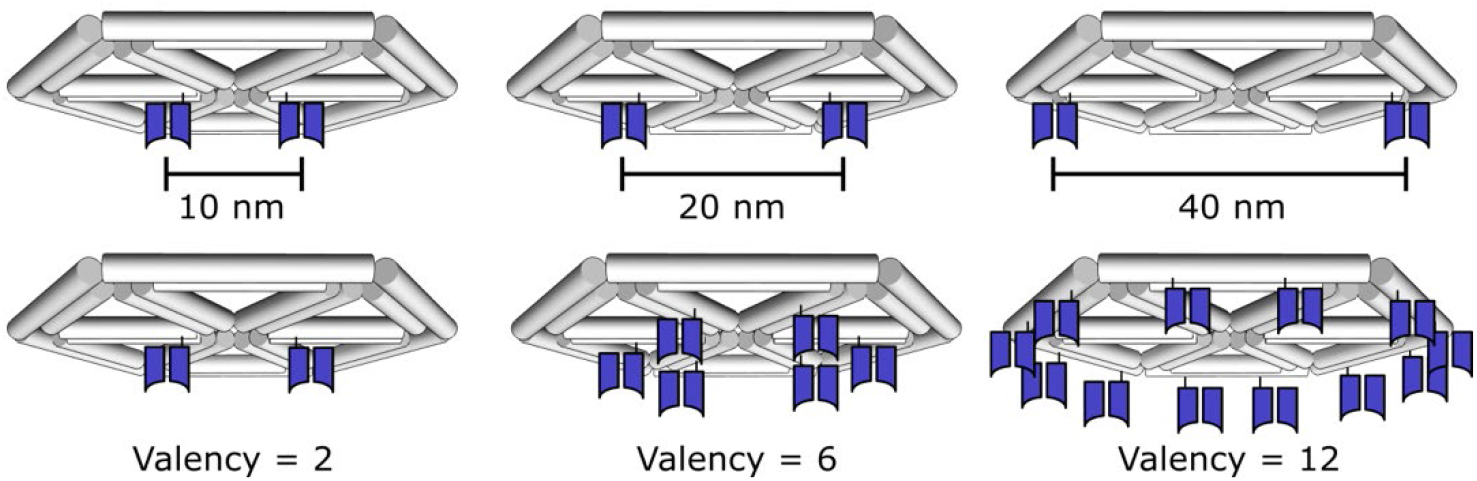
DNA origami nanostructures allow great control over the spatial organization of ligands with nanometre precision and defined composition and number. Double-stranded DNA helices are shown as grey cylinders. Here, ligands are represented as Fab fragments, shown in dark blue.

There are few agonistic antibodies in preclinical development or in the clinic that can induce receptor super-clustering. Receptor clustering can be defined as bringing multiple receptors close together on a 2D membrane to enable activation, while receptor super-clustering brings preformed receptor clusters together, which can increase the agonistic effects even further (**Figure 3**). Similar to other TNFRSF ligands (TNFL), TRAIL forms homomeric trimers.[11] TNFL trimers bind to their corresponding monomeric receptor on a cell surface to initiate trimerization of the receptors to activate downstream signaling pathways.[6] In this context, trimerization of the receptors can be considered as the receptor clustering. Proteins can be engineered to assist receptor clustering, for example single chain mutant variants of TRAIL (SC-TRAIL) were engineered to improve the agonistic effects.[29] This was achieved by linking three extracellular domains of TRAIL by peptide linkers. Here, we combine protein engineering of SC-TRAIL with DNA nanotechnology to super-cluster DR5 and assess cell killing. The methodology described herein could enable control over the spacing of DR5, with defined valencies and geometries, and the ability to study the optimal intraligand distancing. Knowledge of these parameters may enable strong receptor stimulation even at very low concentrations.

**Figure 3.**
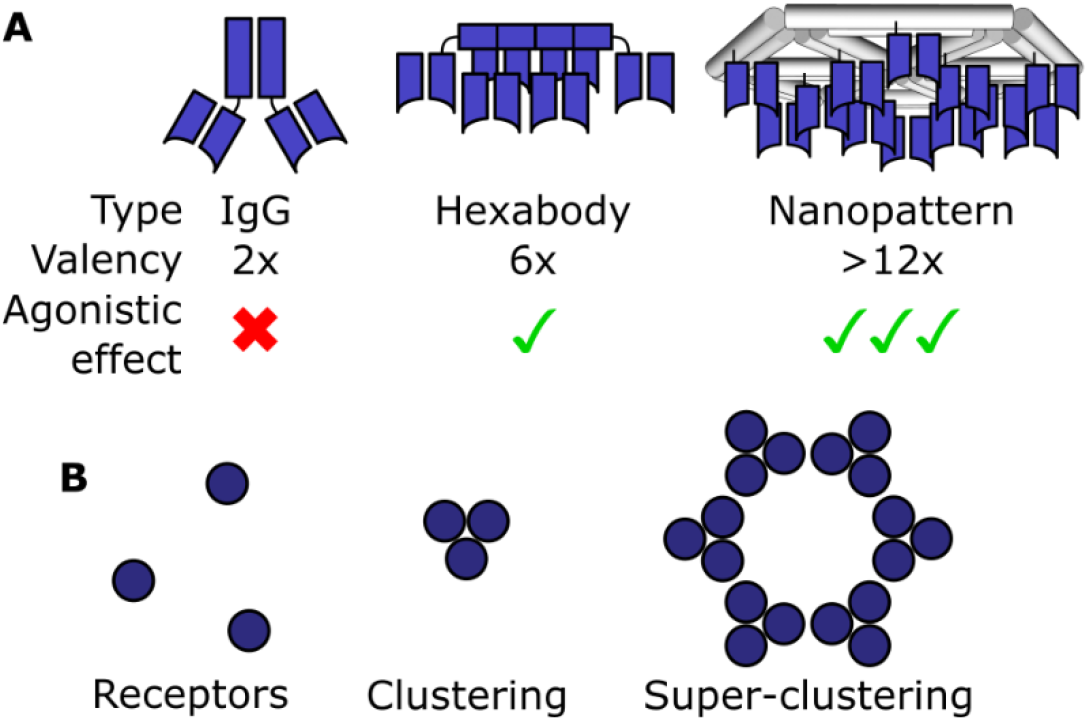
Clustering of receptors improves downstream agonistic stimulation. **A)** DNA nanopatterned antibody/ligand clustering can provide higher binding due to higher avidity, potentially increasing the agonistic effects compared to other super-clustering methods. **B)** Schematic representation of receptor clustering and super-clustering.

## 2 Materials

### 2.1 Reagents and materials

1. M13mp18 genome, Tebubio
2. Amicon Ultra 0.5 – 100 kDa filters (Millipore)
3. Amicon Ultra-15 50 kDa concentrator (Millipore)
4. glow-discharged C-flat carbon film 200 mesh grids (Aurion, Netherlands)
5. 2% aqueous uranyl formate (SPI-Chem, 02545-AA)
6. pCPF3.05 vector: a gift from Robbert Kim (Protein Facility, LUMC)
7. E. coli Rosetta2 (DE3) BL21 derivative (Novagen)
8. HisPur™ Ni-NTA Resin (Thermofisher)
9. Bolt™ 4 to 12%, Bis-Tris, 1.0 mm SDS-PAGE gels (Thermofisher)
10. SimplyBlue™ SafeStain (Thermofisher)
11. HiLoad® 16/600 Superdex® 75 size exclusion chromatography column (Merck)
12. Dibenzocyclooctyne-N-hydroxysuccinimidyl ester (Sigma Aldrich, 761524)
13. Amino modified DNA handles (IDT)
14. Jurkat T cells: a kind gift from Mirjam Heemskerk.

### 2.2 Buffers

1. Folding buffer: 20 mM Tris, 5 mM NaCl and a range of MgCl_2_ concentrations, between 12 to 40 mM, to determine the optimum MgCl_2_ concentration.
2. Nickel binding buffer: 50 mM Tris pH 7.5, 150 mM NaCl and 100 µM ZnCl_2._
3. Wash buffer: 50 mM Tris pH 7.5, 150 mM NaCl, 5 mM MgCl_2_, 100 µM ZnCl_2_, 10 mM imidazole and 10% glycerol.
4. Elution buffer: 50 mM Tris pH 7.5, 150 mM NaCl, 100 µM ZnCl_2_, 250 mM imidazole and 10% glycerol.
5. SC-TRAIL stock buffer: 50 mM Tris pH 7.5, 150 mM NaCl, 100 µM ZnCl_2_ and 10% glycerol.
6. Sortase buffer: 50 mM Tris pH 7.5, 150 mM NaCl, 10 mM CaCl_2_ and 10% glycerol.

### 2.3 Equipment

1. Tecnai T12 Biotwin transmission electron microscope
2. Sonics Vibra-Cellä Ultrasonic Liquid Processor VCX 130
3. ÄKTA™ pure micro FPLC system (Cytiva)
4. TC20 Automated Cell Counter

### 2.4 Software

1. University of California, San Francisco (UCSF) Chimera: https://www.cgl.ucsf.edu/chimera/download.html
2. UCSF ChimeraX: https://www.cgl.ucsf.edu/chimerax/download.html
3. Cadnano2: https://cadnano.org/

## 3 Methods

### 3.1 Determining protein-protein distances to enforce using DNA nanotechnology

1. We manually checked the crystallographic packing of a range of TNFL/TNFR complexes to identify those closely packed in native-like conformations (i.e., all extracellular domains oriented parallel to each other). We identified a structure of a 1BBL/4-1BB complex that conformed to these constraints.
2. We hypothesised that the structurally-homologous TNFL/TNFR family will all allow a similar packing regime, that this is the closest packing that a TNFL/TNFR array could achieve on a cell surface, and that the highest ligand/receptor density may also have the maximum receptor activation.
3. The crystallography data of the 4-1BBL/4-1BB complex can be visualized with UCSF ChimeraX[30] using the command ‘open 6MGP’, which downloads and opens a protein database (pdb) file of a trimer of 4-1BBL molecules, each bound to a 4-1BB receptor (**Figure 4A**)[31].
4. Typing the command ‘unitcell’ shows a small region of the crystal packing of these trimers, and within the 8 copies of the crystallographic unit cell created, there exists a pair of TNF/TNFR trimers closely packed (**Figure 4B**; more easily visible with command ‘del #1 #2.1-4 #2.6 #2.8’).
5. You can see the approximate distance between two of the trimers with the command ‘sel #2.5/C:242@CB #2.7/C:242@CB; distance sel’, which gives a value of ∼5.5 nm.
6. This dimer of trimers can be used to create a hexameric platform, by duplicating the pdb file and repeating the distance and angle of the dimers using the ‘matchmaker’ command (**Figure 4C**).
7. We can take this distance and hexagonal lattice and apply it to DR5/TRAIL to check that there are no steric clashes. We next aligned a model of DR5/TRAIL (pdb code 1D0G [32]) to this hexamer using the matchmaker command, and this model was saved in .pdb format for later use.

**Figure 4.**
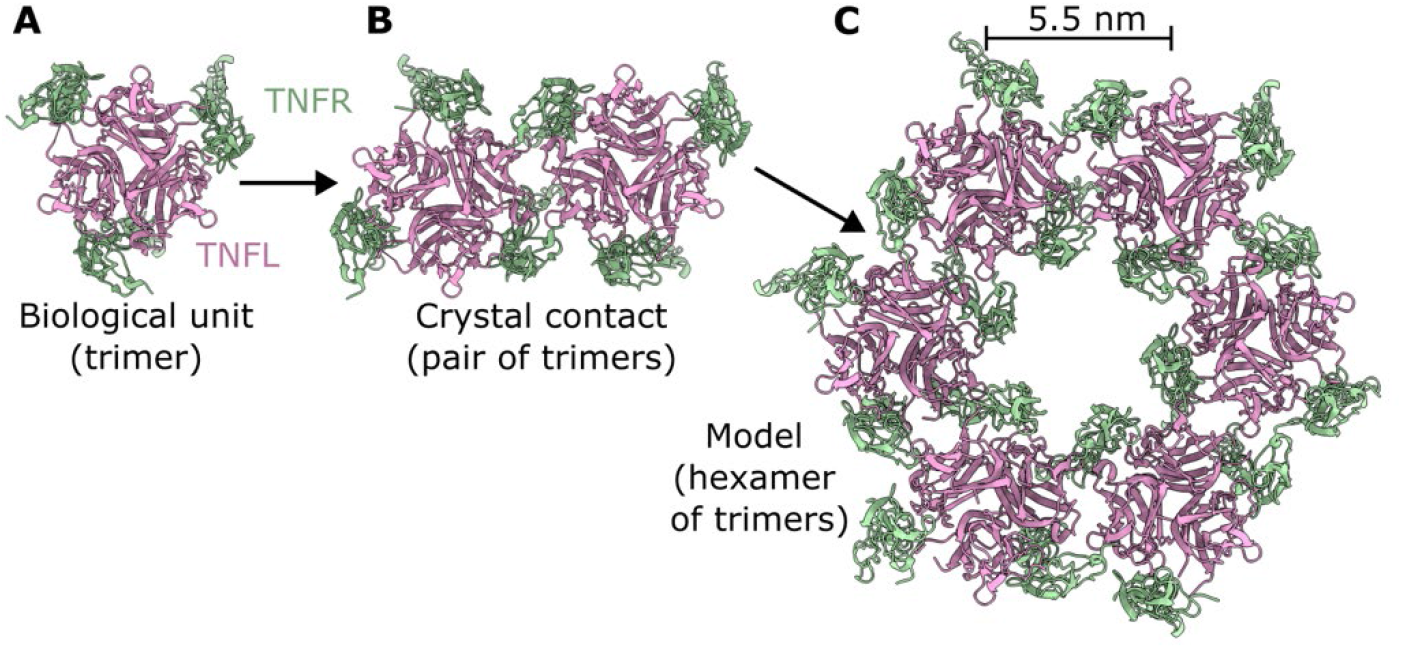
Building a hexameric TNFR model. **A)** Representation of the determination of the intraligand distancing between two different TNFL (pink) trimers when bound to their cognate TNFR complexes (green). Also shown are a dimer of trimers forming crystal contacts **(B)**, and a theoretical model of a hexameric DR5/TRAIL complex **(C)**.

### 3.2 Design a DNA nanostructure

1. The DNA origami nanostructure was designed by writing a .ply file containing the geometry of our DNA design. The files can be created in a number of ways, see refs [33-35]. Below is the ply file for the hexagonal-base pyramid design we use, the visualization of the ply file is shown in **Figure 5A**.

~~~
ply
format ascii 1.0
element vertex 8
property float32 x
property float32 y
property float32 z
element face 12
property list uint8 int32 vertex_indices
end_header
0 0 0
-1 0 0
-0.5 0.866025403784439 0
0.5 0.866025403784439 0
1 0 0
0.5 -0.866025403784439 0
-0.5 -0.866025403784439 0
0 0 1.5
3 0 1 2
3 0 2 3
3 0 3 4
3 0 4 5
3 0 5 6
3 0 6 1
3 7 2 1
3 7 3 2
3 7 4 3
3 7 5 4
3 7 6 5
3 7 1 6
~~~
2. The ply file can be loaded into Athena[33] or alternatively can be uploaded onto one of the servers from the Bathe lab[34,36,37] to create all the required files.
3. In Athena, select Talos to create nanostructures with 6 DNA helices per edge (**Figure 5B**), which are more rigid than the alternative Daedalus designs that use only 2 DNA helices per edge length.[35]
4. Set the desired side length; 42 base pairs in this case, and then “Run Sequence Design”. Next, select “Include PDB” and press Save. Athena will create all the files required to modify and synthesise a DNA nanostructure with the geometry provided in the ply file, including an atomic model (in pdb format; **Figure 5C**) and an associated cadnano file (in json format; **Figure 5D**)

**Figure 5.**
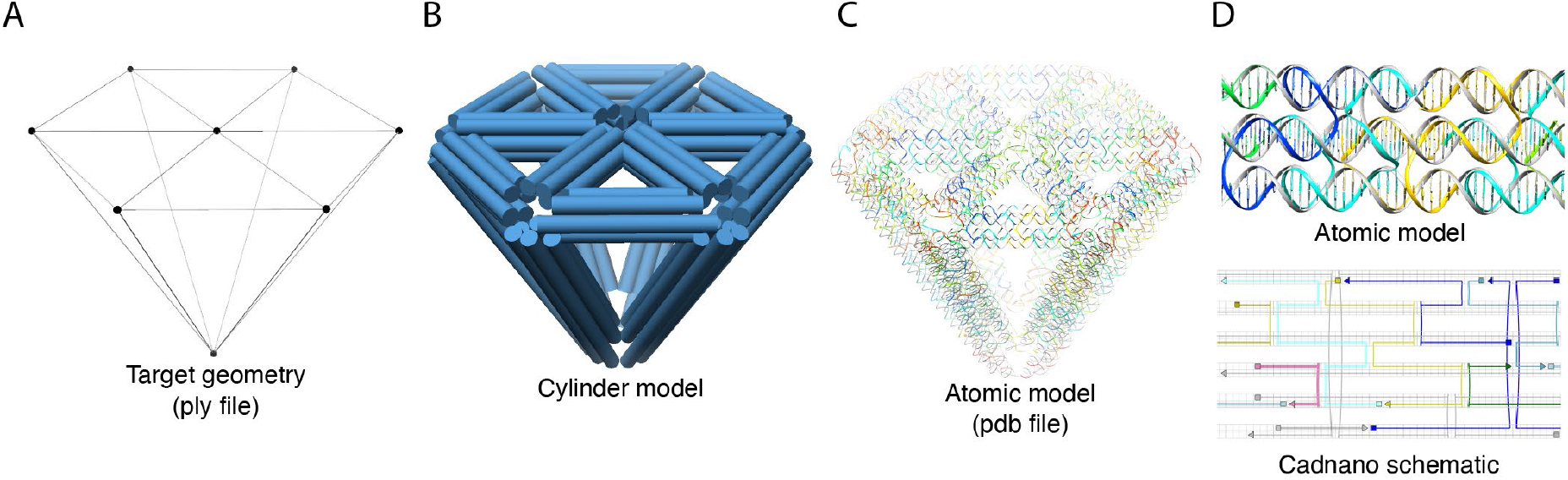
Creation and modification of the DNA nanostructure using Athena.

### 3.3 Identify locations on the DNA nanostructure to correctly place ligands

1. Specific staple strands within the DNA nanostructure will be modified to bind to proteins; it is the location of these staple strands within our DNA nanostructure that will determine the spacing of the proteins. To determine which staples to modify, the atomic model obtained from Athena[33] was opened in UCSF Chimera[38], together with the TRAIL/DR5 hexamer model that we created earlier in step 1. **Figure 6A** shows a simplified model with white and blue representing the scaffold and staple strands of the DNA origami nanostructure, respectively, and pink the TRAIL proteins. The two structures have been moved with respect to one another such that the proteins are all above DNA helices.
2. In **Figure 6B**, the staple strands close to the termini of the TRAIL proteins have been coloured red; these are the staples that can be modified to present the proteins on a hexagonal array with a distancing of ∼5.5 nm.
3. Also open the *guide_caDNAno.bild file output by Athena in UCSF Chimera (**Figure 6B**) (see Note 1). The numbers of the DNA helices that contain the red staple strands can be seen, and these correspond to the numbers of the helices in the cadnano .json file (explained below).
4. The specific model IDs for the red staples were noted with the corresponding DNA sequence, which can be obtained with the sequence tool in UCSF Chimera (Tools > Sequence > Show Sequence Viewer) (**Figure 6C**).

**Figure 6.**
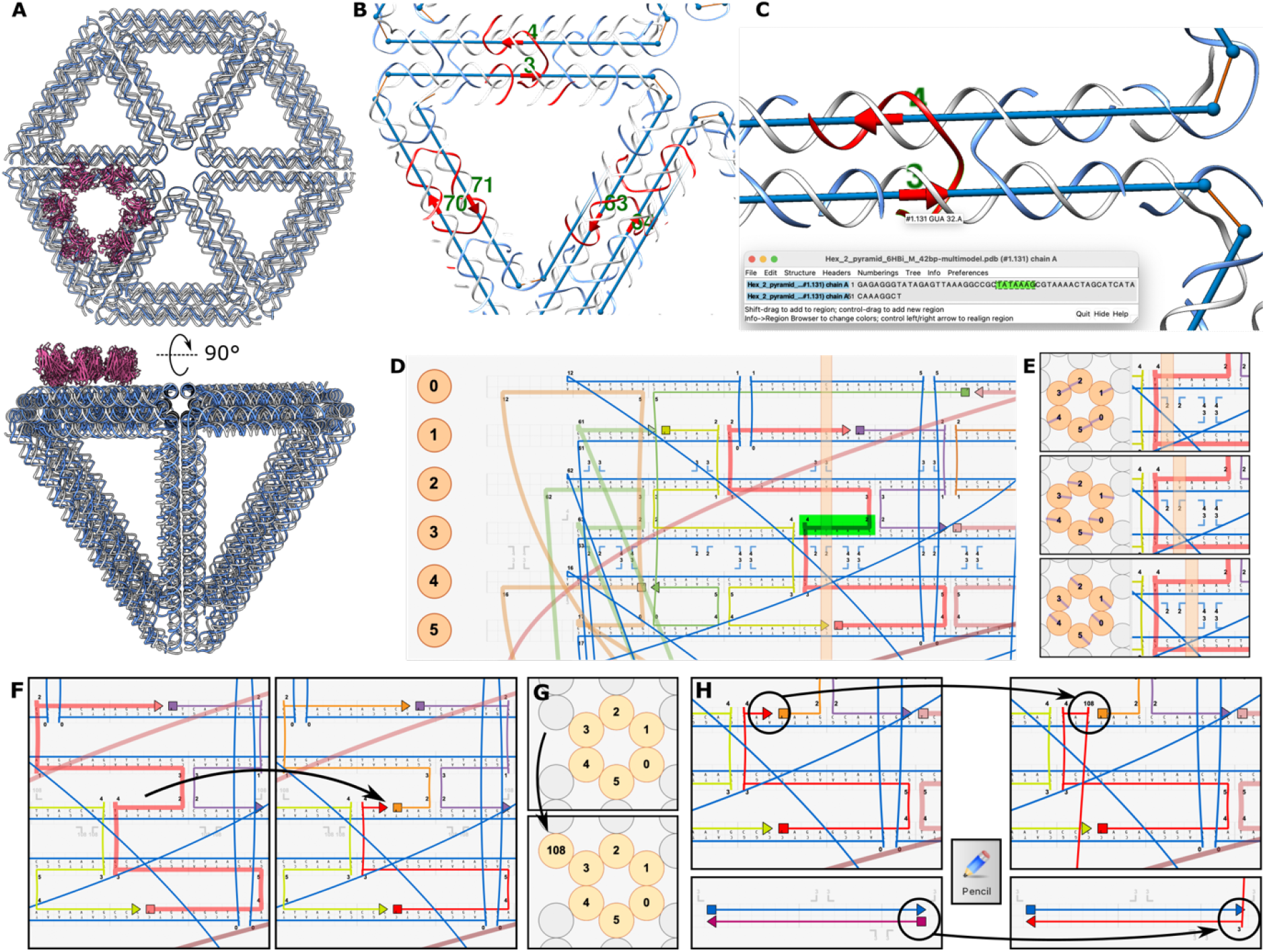
Workflow of modifying the staple DNAs overlapping with the hexamer-of-trimers TRAIL model. **A)** Models of the DNA nanostructure and hexameric TRAIL complex aligned so that each trimer is above a DNA helix. **B)** Strands in locations that could be modified are coloured red, then the protein model is hidden for convenience. **C)** The ID of the helices is determined in UCSF Chimera and the sequence shown. **D)** The ID of the helices is shown in Cadnano, and the sequences matched to those of the pdb model to confirm the correct position is identified. **E)** The pitch of the DNA helices is visible as blue lines within the orange circles, which represent DNA helices. As the orange bar is moved right, the blue line rotates clockwise, indicating the position of the scaffold strand. The staples are oriented directly opposite this blue line. The middle panel shows the situation where the staple strand is pointing directly out from the DNA nanostructure. **F)** At this location, the staple can be broken in order to modify it. **G)** An additional empty DNA helix is added at a convenient location (it does not need to be in any particular location; here it is shown adjacent to strand 3 for convenience). **H)** A new DNA helix of the desired length is added to the new empty helix, before being connected to the previously-identified location. Adding the DNA handle is now complete.

### 3.4 Identify locations in the Cadnano file to correctly place ligands

1. Open the .json file in Cadnano[39], where each double-stranded DNA helix is numbered (represented by the orange circles in **Figure 6D**), and these correspond to the numbering in the “*guide_caDNAno.bild” file output from Athena.
2. The sequence of the scaffold strand can be added in Cadnano to further aid identification of each staple strand.
3. By comparing these files – the pdb model, the *guide_caDNAno.bild file, and the Cadnano[39] file – the locations identified in UCSF Chimera can be identified to determine which staples to modify. For example, in **Figure 6C & D**, the same sequence and location has been highlighted in green.
4. In **Figure 6E**, the orange circles represent the DNA helices. Within the orange circles are blue lines, which show the pitch of the scaffold DNA at the position shown by the orange rectangle on top of the DNA map. As the rectangle is moved right, the blue lines rotate clockwise (DNA is right-handed). Importantly, the staple DNA strands are at the opposite side to the scaffold strand, and therefore opposite to the blue lines.
5. We want our strands to be modified pointing away from the DNA nanostructure. Therefore, to modify helix 3, we have chosen the location of the middle panel in **Figure 6E** to include an extension to the staple, where the blue line points directly *in* to the DNA nanostructure, and the staple therefore points directly *out* of the nanostrucutre (this location can be confirmed by comparing to the pdb model), and will therefore correctly display the protein. Thus, we will break the staple at the indicated position (**Figure 6F**). (see note 2)

### 3.5 Modify staple strands on the DNA nanostructure for ligand binding

1. After locating the positions of the modifications, we need to extend the strands by 18-20 nucleotides. To do this, create an empty DNA helix by placing a new circle next to the strand you wish to modify (**Figure 6G**). In this case, the strand is number 108, but this depends on the number of helices in your DNA nanostructure.
2. In the row for strand 108, draw a new scaffold/staple pair of the same length as the extension, in this case 19 nucleotides long (**Figure 6H**).
3. Finally, using the pencil tool, connect the relevant termini of the staple strand and the extension (**Figure 6H**). The desired sequence of the extensions can also be added at this point.
4. In this example, we have indicated 6 positions to modify the staple strands, but this DNA nanostructure can accommodate 24 modifications on a 5.5 nm lattice.
5. When all extensions have been added at the desired locations, it is helpful to colour all the staples as groups. We usually use at least two colours; “core” staples – those that form the main structure – and “modification” staples – those that will display the proteins.
6. The sequence file can be exported from Cadnano as a .csv file. There will be numerous question marks ‘?’ in the sequence file; these correspond to the unpaired single-stranded DNA staples that link the structure together.[34] These can all be replaced as Thymine (T) bases. In our case, all of the staples were modified to not exceed the 60-nucleotide limit including the extension sequence, as this saves a significant amount of money when ordering the staples.
7. The staple sequences were then ordered from Integrated DNA Technologies (IDT), already dissolved in water at a concentration of 100 µM, to be used for the folding reactions.

### 3.6 Design, folding and purification of DNA origami nanostructures

1. After receiving the staples from IDT, the staples were thawed overnight at 4°C.
2. The single stranded viral M13mp18 genome was as the scaffold. The final concentration of the scaffold DNA and each of the oligonucleotide strands from IDT were 20 nM and 200 nM in the folding reactions mixture, respectively.
3. After mixing together all required components in folding buffer, the folding reactions were placed in a Bio-Rad C1000 Touch Thermal Cycler and thermally annealed. To do so, the mixtures were held at 80°C for 1 minute, followed by a thermal annealing ramp from 80°C to 75°C (at a rate of 0.2°C/min), then subsequently from 75°C to 30°C (0.1°C/min) and finally from 30°C to 20°C (0.1°C/min). **Figure 7A** shows the agarose gel analysis of this folding reaction. The small shift in bands represents the folding of the scaffold DNA. The intensity of the bands represents the amount of folded DNA nanostructures.
4. It can be observed that the intensity of the bands decreases for MgCl_2_ concentrations higher than 28 mM, suggesting that the folding reaction efficiency decreases above this concentration of MgCl_2_. The excess staple DNA pool is also visible.
5. After determining that ∼20 mM MgCl_2_ was optimal, we then prepared more folding reaction, this time with a range of NaCl concentrations, from 0 to 320 mM, to determine the optimum salt concentration (data not shown).[28]
6. The folded structures were purified from excess oligonucleotides using the Amicon Ultra 0.5 – 100 kDa filters by washing five times with the folding buffer containing the determined optimum salt concentrations.

**Figure 7.**
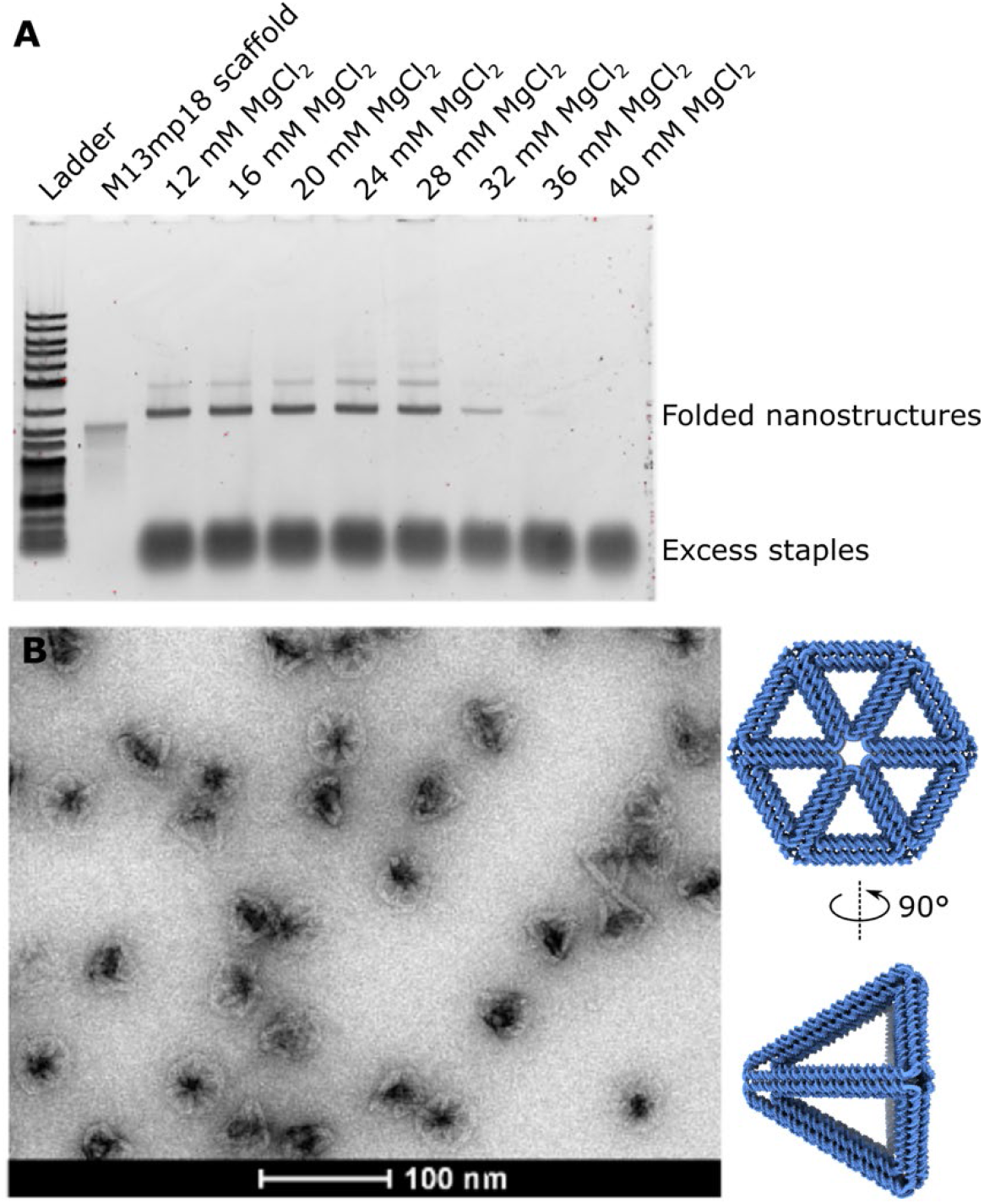
Folding and characterisation of DNA origami nanostructure. **A)** Agarose gel analysis of the folding reactions with various MgCl_2_ concentrations to determine the optimum salt concentrations. The locations of both excess staples and folded DNA origami nanostructures are indicated. 1% agarose gel was run at 100 V for 1 hour with 3 µL samples that were diluted 12.5x in MQ. **B)** Schematic representation of the designed DNA origami nanostructure with the corresponding transmission electron microscopy images obtained with negative staining. Scale bar: 100 nm.

### 3.7 Electron microscopy of folded DNA nanostructures

1. We prepared negative staining grids of the folding reaction with 20 mM MgCl_2_ to inspect the folded nanostructures with transmission electron microscopy (TEM).
2. To do so, 10 µL of purified or unpurified samples were applied at 2 nM to glow-discharged C-flat carbon film 200 mesh grids and incubated for 1 minute.
3. After removal of the excess solution, staining was performed using 10 µL of 2% uranyl formate and incubated for 10 seconds.
4. After removal of the solution, the grid was air dried for 5-15 minutes.
5. Images were obtained using a Tecnai T12 Biotwin transmission electron microscope operated at a voltage of 120 kV. **Figure 7B** shows one of the representative images with correctly folded DNA nanostructures.

### 3.8 Protein design to covalently link SC-TRAIL to DNA

1. Various drugs or materials can be loaded to functionalize the DNA nanostructures (see note 3). We recommend using DNA hybridization or specific interactions between nanobodies and their small targets for functionalization.[40]
2. Specifically, we use a sortase enzyme[41] to covalently link proteins directly to DNA oligonucleotides, then hybridize these to the folded DNA nanostructure. This requires expressing SC-TRAIL with a with C-terminal sortag – a short peptide sequence LPXTG (where X is any amino acid – we use E) – followed by a 6-His tag for purification.
3. The gene for SC-TRAIL therefore comprised three concatenated repeats of TRAIL with 3× gly-gly-ser linkers between each monomer, with a C-terminal sortag and 6-His tag:

~~~
MGPQRVAAHITGTRGRSNTLSSPNSKNEKALGRKINSWESSRSGHSFLSNLHLRNGELVI
HEKGFYYIYSQTYFRFQEEIKENTKNDKQMVQYIYKYTSYPDPILLMKSARNSCWSKDAE
YGLYSIYQGGIFELKENDRIFVSVTNEHLIDMDHEASFFGAFLVGGGSGGSGGSPQRVAA
HITGTRGRSNTLSSPNSKNEKALGRKINSWESSRSGHSFLSNLHLRNGELVIHEKGFYYI
YSQTYFRFQEEIKENTKNDKQMVQYIYKYTSYPDPILLMKSARNSCWSKDAEYGLYSIYQ
GGIFELKENDRIFVSVTNEHLIDMDHEASFFGAFLVGGGSGGSGGSPQRVAAHITGTRGR
SNTLSSPNSKNEKALGRKINSWESSRSGHSFLSNLHLRNGELVIHEKGFYYIYSQTYFRF
QEEIKENTKNDKQMVQYIYKYTSYPDPILLMKSARNSCWSKDAEYGLYSIYQGGIFELKE
NDRIFVSVTNEHLIDMDHEASFFGAFLVGGGSGGSGGSLPETGGHHHHHH
~~~
4. This gene was ordered from Invitrogen GeneArt and cloned into the pCPF3.05 vector.

### 3.9 Expressing sortagged SC-TRAIL

1. E. coli Rosetta2 (DE3) with pCPF3.05 encoding for SC-TRAIL was first inoculated in 100 mL Erlenmeyers with 10 mL LB-medium containing 50 μg/mL kanamycin in a shaker at 37°C, 200 rpm for 17-18 hours.
2. The culture broth was then inoculated in two 2 L Erlenmeyers containing 500 mL LB-medium with 50 μg/mL kanamycin until an OD 600 value of 0.5-0.6 was reached. 1 mM IPTG was applied to initiate the production of SC-TRAIL.
3. The culture broth was sampled and centrifuged at 6000 g for 1 minute and the pellet was dissolved in 50 μL Laemmli sample buffer (50 mM DTT) before addition of IPTG to monitor the SC-TRAIL production at the starting point.
4. Incubation took place at 18°C 200 rpm for about 24 hours, after which the culture broth was sampled and centrifuged again as described above. The culture was then harvested by centrifugation at 4000 g, 4°C for 20 minutes and the pellet was washed with 10 mL Nickel binding buffer and transferred to a 50 mL tube.
5. The ∼2.5 g pellet was dissolved in 10 mL Wash Buffer. The solution was afterwards sonicated with Sonics Vibra-Cellä Ultrasonic Liquid Processor VCX 130 (total time: 1 min 45 seconds, pulse on: 15 seconds, pulse off: 45 seconds, amplitude: 50%).
6. The supernatant was collected after centrifugation (20 minutes at 16000 g, 4°C).
7. Protein purification was performed with HisPur™ Ni-NTA Resin.
8. The protein was eluted with different concentrations of imidazole containing Elution Buffer ranging from 50 mM – 250 mM imidazole.
9. Samples of the elutions were studied on a Bolt™ 4 to 12%, Bis-Tris, 1.0 mm SDS-PAGE gels with a running scheme of 200 V for 35 minutes.
10. Proteins were visualized by staining with SimplyBlue™ SafeStain.
11. SC-TRAIL containing samples were further purified and buffer exchanged to SC-TRAIL stock buffer using ÄKTA™ pure micro with HiLoad® 16/600 Superdex® 75 pg with 1 ml/min.
12. Elutions containing the SC-TRAIL were then concentrated using 15 mL Amicon Ultra-15 50 kDa (10 minutes, 3000 g, 4°C).

### 3.10 Sortase-mediated transpeptidation reaction

1. A small click handle was attached to the SC-TRAIL using the transpeptidation reaction mediated by the sortase enzyme (**Figure 8**, reaction *i*). This was followed by the click reaction between the azide-containing SC-TRAIL and DBCO containing DNA handles (**Figure 8**, reaction *ii*). Finally, addition of DNA-SC-TRAIL conjugates to the final DNA nanostructures in excess facilitates functionalisation of the nanostructure (**Figure 8**, reaction *iii*)
2. Conjugation was performed by incubating 50 µM SC-TRAIL, 1 mM functional peptides containing FITC (as a positive control) or azide, 25 µM sortase 5M, in sortase buffer. Martijn Verdoes (Radboud University) kindly gifted the plasmid encoding for Sortase 5M.
3. The sequence of the functional peptide was NH_2_-GGG-K(X)-G-H, where X is the FITC fluorophore or N_3_ azide on the lysine sidechain.
4. The reaction mixture was incubated at 4°C, mixing for total of 2 hours, followed by centrifugation using Amicon Ultra-0.5 50 kDa (10 minutes, 3000 g, 4°C) to remove the sortase enzyme and the excess peptides.
5. The reaction was monitored with SDS-PAGE Bolt™ 4 to 12%, Bis-Tris, 1.0 mm from Thermo Fisher Scientific, with a running scheme of 200 V for 35 minutes.
6. Proteins were stained with SimplyBlue™ SafeStain from Thermo Fisher Scientific (**Figure 9A**), whilst the fluorescent signal from the FITC fluorophore could be measured with excitation and emission at 490 nm and 525 nm, respectively (see note 4). Overlap of the fluorescent bands with the TRAIL bands confirmed the successful modification (**Figure 9A**).
.
7. Separately, an azide containing peptide was conjugated to the SC-TRAILs, which will be used to conjugate a small oligo to allow functionalization of the DNA nanostructure with the ligands.

**Figure 8.**
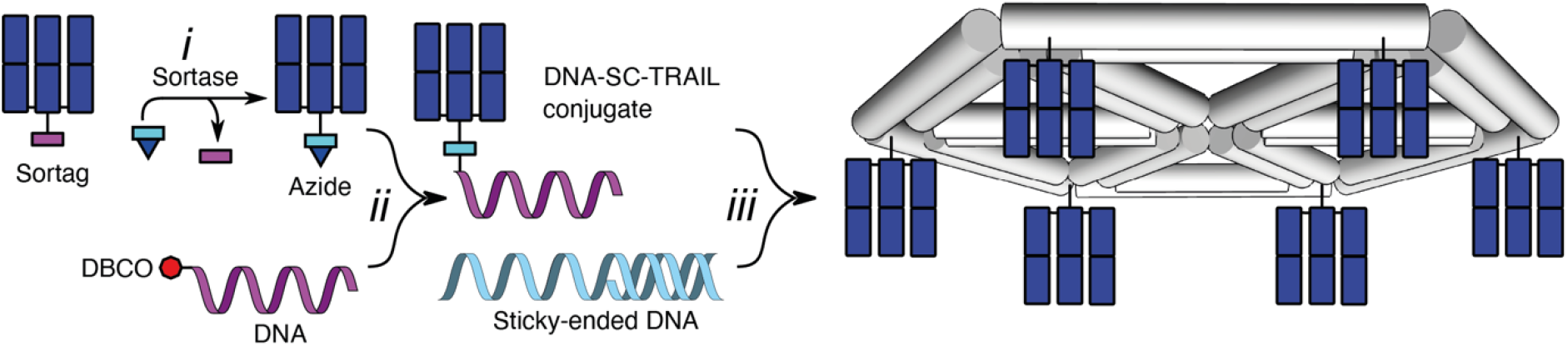
Scheme for incorporating SC-TRAILs into DNA nano-structures using sortase reactions and click chemistry. Reaction **i** describes the usage of sortase enzyme for transpeptidation reaction for addition of azide to the protein. Reaction **ii** describes the click-reaction for covalently attachment of the DNA handle to the SC-TRAILs. Reaction **iii** shows the usage of DNA hybridisation for the functionalisation of DNA nanostructures.

**Figure 9.**
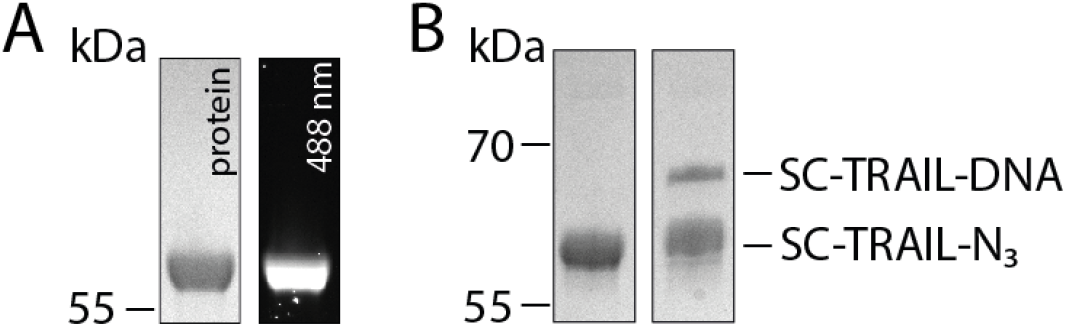
Conjugation of TRAIL proteins with functional groups. **A)** SDS PAGE analysis of SC-TRAIL conjugation with a fluorescent peptide, GGG-K(FITC)-G. Overlap of fluorescent bands with SC-TRAIL bands confirms the introduction of the fluorescent peptide. **B)** SDS PAGE analysis of SC-TRAIL conjugation with the DNA handle. The shift in the band size when the reaction is performed confirms the conjugation of the DNA handle. The band on the left represents the SC-TRAIL-sortag, the band on the right bottom represents SC-TRAIL-N_3_ and the band on the right top represents the conjugation product SC-TRAIL-DNA.

### 3.11 Click reaction to link SC-TRAIL to DNA

1. 1.5 mg of Dibenzocyclooctyne-N-hydroxysuccinimidyl (DBCO) ester was weighed and dissolved in 300 µL DMSO and this solution was further used to prepare DBCO-functionalized DNA handles by mixing 20 µL DBCO ester, 20 µL DMSO, 20 µL PBS and 20 µL amino modified DNA handles.
2. The mixture was incubated overnight at 37°C, 1100 rpm.
3. DBCO-modified DNA handles were then purified with Amicon Ultra-0.5 10 kDa (10 minutes, 13000 g, RT) and the concentration was determined with the absorbance at 260 nm.
4. SC-TRAIL-N_3_ was incubated with the DBCO-functionalized DNA handle at 4°C overnight with 3 µM ligand and 30 µM oligo.
5. SC-TRAIL-DNA was then purified from the excess DNA-DBCO using Amicon Ultra-0.5 50 kDa (10 minutes, 3000 g, 4°C).
6. The reaction was analyzed with an SDS-PAGE to monitor the reaction (**Figure 9B**); the shift of the band confirmed successful labeling of the ligands.
7. The ratio between the SC-TRAIL-N_3_ and SC-TRAIL-DNA was estimated using the SDS-PAGE and this ratio was later used to estimate the concentration of SC-TRAIL-DNA after determination of the concentration with absorbance at 280 nm.

### 3.12 Functionalization of DNA nanostructures with SC-TRAIL

1. The DNA-origami nanostructures (50 nM) were incubated overnight at room temperature with 24× molar concentration of SC-TRAIL-DNA (as there are 24 binding sites).
2. This was followed by purification with Amicon Ultra-0.5 100 kDa filters (4 minutes, 3000 g, RT).
3. The final concentration was determined by absorbance at 260 nm.

### 3.13 Cell killing induced by DNA-templated SC-TRAIL

1. Jurkat cells were seeded 100 x 10^3^ cells/well in a 96-well plate in 45 or 20 µL overnight at 37°C, 5% CO_2_.
2. Cells were treated with different concentrations of SC-TRAIL containing DNA origami nanostructures and incubated at 37°C, 5% CO_2_ overnight.
3. Viable cells were counted next day using TC20 Automated Cell Counter from Bio-Rad with trypan blue staining.
4. **Figure 10** shows the cell killing properties of the constructs with the SC-TRAIL on TRAIL-sensitive Jurkat cells. For this demonstration we compared the cell killing of an empty DNA origami construct (red Ori-0; no protein, 2 nM DNA concentration) and a construct with 24 SC-TRAIL binding positions (green Ori-24; 8 and 16 nM SC-TRAIL on 0.33 and 0.66 nM DNA concentration, respectively) at two different concentrations. As **Figure 10** shows, empty DNA origami construct does not induce any cell killing compared to the media control. However, the number of live cells decreases drastically when the construct with Ori-24 is used. Interestingly, SC-TRAIL-sortag alone (purple) does not induce efficient cell killing. This therefore demonstrates the additional efficacy resulting from TRAIL-induced DR5 super-clustering using protein engineering and DNA-mediated nanopatterning.

**Figure 10.**
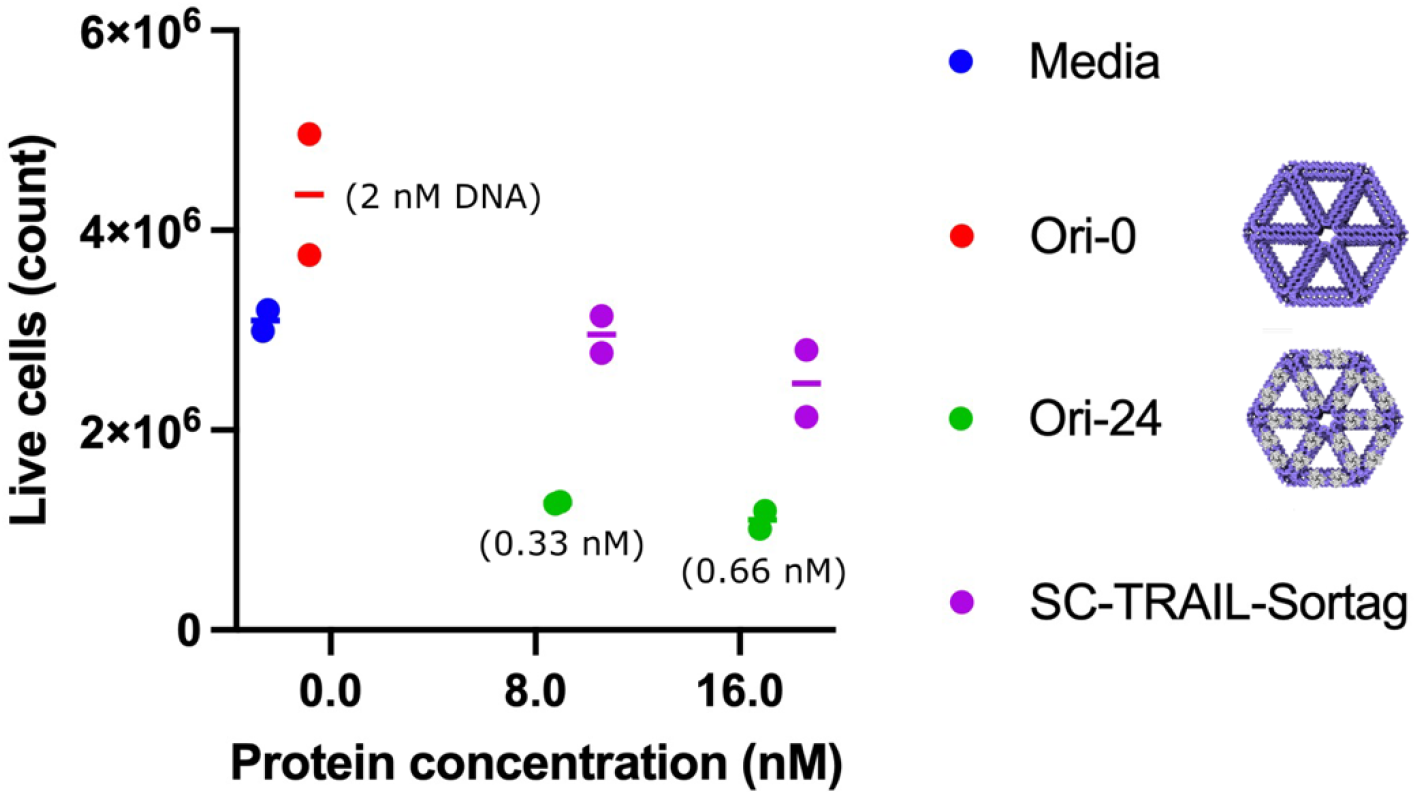
Super-clustering SC-TRAIL using DNA-nanotechnology results in enhanced cell killing. Live cell counts of Jurkat cells treated with DNA origami constructs containing SC-TRAIL or SC-TRAIL-sortag at different concentrations. DNA concentrations are shown in parentheses.

## 4 Notes

1. Use UCSF Chimera to open these, do not use UCSF ChimeraX, as this will not display the numbering correctly.
2. There are other important parameters to be aware of, including the length of the staple strand (ideally 20-60 bases long), the proximity of adjacent Holliday junctions (not too close, not too far), and the directionality (i.e., whether to extend the 3’ or 5’ of the DNA). We do not cover these considerations here, and we instead refer to reference [28] for more information.
3. Different approaches can be used to functionalize the DNA nanostructures, for example noncovalent binding by DNA hybridization,[25] covalent binding by click chemistry,[42] using the interaction between biotin and streptavidin[43] or intercalation ability of doxorubicin can be exploited to load drugs,[44,45] also reviewed in [46]. Oftentimes, because of its ease in use and ready availability, the interaction between streptavidin and biotin is used to functionalize DNA nanostructures. However, we avoid using streptavidin for functionalization, because of the fact that streptavidin has four binding domains for biotin and can aggregate DNA nanostructures by binding to multiple biotins from multiple DNA nanostructures.
4. The fluorescent signal has to be measured before the staining with SimplyBlue™ SafeStain to visualize the signal. Oftentimes, the fluorescent signal from the excessive FITC-peptide saturates the fluorescent signal, to solve this problem, lower part of the gel can be cut to remove the fluorescent peptides.

